# Core microRNAs regulate neural crest delamination and condensation in the developing trigeminal ganglion

**DOI:** 10.1101/2025.06.27.661967

**Authors:** Rocío B. Marquez, Estefanía Sánchez Vázquez, Andrés M. Alonso, Yanel E Bernardi, Emilio M. Santillan, Peter Lwigale, Luisa Cochella, Marianne E. Bronner, Pablo H. Strobl-Mazzulla

## Abstract

Cranial neural crest cells (NCCs) undergo dynamic processes during embryonic development, including delamination from the neural tube by epithelial-to-mesenchymal transition (EMT), migration to the periphery, condensation via mesenchymal-to-epithelial transition (MET) and differentiation into structures like the trigeminal ganglion. Here, we identify and characterize the function of a core set of miRNAs involved in these transitions during the formation of the trigeminal ganglion in the chick embryo. We further identify putative targets of miRNAs involved in neural crest EMT and MET. Notably, introducing MET-involved miRNAs into trunk NCCs endows these cells with the ability to condense and differentiate into neurons *in vivo* in a manner reminiscent of cranial rather than trunk NCCs. Our findings shed light on the intricate regulatory networks governing NCC behavior, positioning miRNAs as key regulatory elements required for migratory transitions and axial level specific differentiation capabilities.

## INTRODUCTION

Neural crest cells (NCCs) are an embryonic stem cell type present in all vertebrates and characterized by their extensive migratory ability and multipotency ^1^. Initially arising within the forming central nervous system, NCCs subsequently undergo an epithelial-to-mesenchymal transition (EMT) to leave the neuroepithelium and initiate migration to diverse locations throughout the developing embryo. During EMT, pre-migratory NCCs undergo significant changes in their transcriptional repertoire enabling them to transition to a migratory cell type ^2^.

NCCs arise along the entire body but exhibit distinct molecular signatures and differentiation potentials depending upon their axial level of origin. While initially multipotent, they undergo sequential regulatory states before committing to their final fate ^1,3,4^. In the cranial region, a subset of NCCs halts migration and undergoes a condensation process much like the reverse of EMT, termed mesenchymal-to-epithelial transition (MET) ^5,6^. This occurs before they differentiate into neurons and glial cells that form the sensory ganglia of the head ^7,8^ the largest of which is the trigeminal ganglion, a critical component of the peripheral nervous system that relays sensory information such as pain, touch, and temperature from the head and various organs to the brain ^9^.

As shown in the chick embryo, the EMT resulting in NCC delamination from the neural tube involves the loss of Cadherin6b, which is directly down-regulated by a transcriptional repressor complex comprised of SNAIL2 and PHD12 ^10,11^. Additionally, several types of cell-cell and cell-matrix interactions guide cranial and/or trunk NCC migration and condensation to ensure proper gangliogenesis, including Eph-ephrin ^11,12^, neuropilin-Semaphorin ^13^, laminin-extracellular matrix ^14^, Slit1-Robo2 signaling ^15^, and integrin-extracellular matrix ^16^ interactions. Notably, heterotopic transplantation experiments in birds have demonstrated a graded potential of NCCs to form sensory neurons along the rostrocaudal axis ^3^. For example, more caudal neural crest populations like the cardiac neural crest, which arises from the hindbrain, and trunk neural crest, from spinal cord levels, form much smaller trigeminal ganglia with fewer somatosensory neurons when grafted in place of the cranial neural crest. These results suggest that there is intrinsic information in NCCs at different axial levels that may contribute to their potential to condense and generate trigeminal neurons.

The regulatory mechanisms underlying neural crest EMT, followed by migration and reaggregation mediated by MET, likely involve a tightly regulated and evolutionarily conserved gene regulatory network (GRN) ^17,18^. During both morphogenesis and pathological conditions such as cancer metastasis ^19–22^, microRNAs (miRNAs) have been identified as key regulators of EMT and MET transitional states. Moreover, there is growing evidence supporting a role for miRNA-mediated gene silencing in NCC induction and delamination across various animal models, including zebrafish, Xenopus, chicken, and mice ^23–27^. For example, we previously described the function of a single microRNA, miR-203, whose epigenetic repression is required for the initial delamination of NCC ^25^. Later, its re-expression is essential for NCC-placode communication and the condensation process that leads to trigeminal ganglion morphogenesis ^28^. These findings underscore the importance of miRNAs in regulating both the delamination and condensation transitions during NCC development.

While transcription factors have been extensively studied in these processes, the role of microRNAs (miRNAs) remains underexplored, particularly in the context of MET-like events. To obtain a more complete picture of the role of miRNAs during critical stages of cranial neural crest EMT and MET, here we performed miRNA transcriptome profiling on sorted NCCs during pre-migratory, migratory, and condensing stages to identify a core set of miRNAs involved in their delamination (EMT-miRNAs) and condensation (MET-miRNAs) to form the trigeminal ganglion. Using bioinformatics prediction and transcriptome profiling, we identified putative targets and functionally characterized their roles through loss-of-function experiments. Finally, we find that introducing MET-miRNAs into trunk NCCs from GFP-transgenic chick embryos and grafting them into an ectopic cranial environment of host wild-type embryos endowed these cells with cranial crest-like ability to condense and differentiate into neurons of the trigeminal ganglia. These findings underscore the importance of miRNAs as mediators of epithelial plasticity and intrinsic axial level-specific regulators of neural crest developmental potential.

## MATERIALS AND METHODS

### Embryo Preparation and Electroporation

Fertilized White Leghorn chick embryos were obtained from “Escuela de Educación Secundaria Agraria de Chascomús” (Chascomús, Argentina) and incubated at 38° C until the embryos reached the proper developmental stage. Chicken embryos were collected and cultured according to the Early Chick (EC) protocol and staged based on (HH) ^29^ or by counting the number of somite pairs (somite stage, ss). For *ex ovo* electroporation: Chicken embryos at stage HH4-5 were transfected as previously described ^30,31^. Plasmids, miRNA inhibitors or mimics were injected by air pressure using a glass micropipette targeting the epiblast of embryos collected on filter paper rings. Platinum electrodes were placed vertically across the chick embryos and electroporated with five pulses of 5.1 V in 50 ms at 100 ms intervals.

For *in ovo* electroporation: Plasmids, miRNA inhibitors or mimics were injected by using a glass micropipette with air pressure. Platinum electrodes were positioned horizontally adjacent to the developing neural tube (5 15.5 V at 50 ms at 100 ms intervals). Penicillin and streptomycin (100 g/mL) were added to the Ringer solution to prevent contamination. The electroporated eggs were then sealed with adhesive tape and incubated at 38°C to reach the desired stage ^32^. In all treatments the embryos were injected only into the right side of the embryo and phenotype comparisons were made comparing the untreated side. Embryos with weak and/or patchy electroporation or with strong morphological abnormalities were discarded.

### Cell Dissociation and FACS Sorting

To isolate pre-migratory and migratory NCC, embryos were *ex ovo* electroporated with the NCC-specific Foxd3 enhancer NC1 ^33^, driving expression of a Citrine fluorescent reporter (2 mg/ml). For condensing NCC, embryos were *in ovo* electroporated with NCC-specific Sox10 enhancer 10E2 ^34^. GFP-positive (NCCs) and GFP-negative (Control) cells were collected via fluorescence-activated cell sorting (FACS) at three key stages of cranial NCC development: HH9 (pre-migratory), HH10 (migratory) and HH16 (condensing). Cranial regions and trigeminal ganglion from electroporated embryos were microdissected and enzymatically dissociated using 1.5 mg/mL dispase in DMEM supplemented with 10 mM HEPES (pH 7.5) at 37°C for 15 minutes, with gentle pipetting to facilitate single-cell suspension. A final dissociation step was performed using 0.05% trypsin at 37°C for 3 minutes. The enzymatic reaction was stopped, and cells were re-suspended in Hanks buffer (1X HBSS, 0.25% BSA, 10 mM HEPES, pH 8). Following centrifugation at 500 x g for 10 minutes, cells were resuspended in fresh Hanks buffer, filtered through 40 μm strainers, and centrifuged again at 750 × g for 10 minutes. The resulting cell pellet was resuspended in 500 μL of Hanks buffer. GFP-positive cells were sorted using a BD FACS-Aria Fusion cell sorter. On average, approximately 3000 purified NCC were subsequently used for both bulk miRNA-seq profiling analyses.

### Small RNA Extraction, Library Preparation, and Sequencing

FACS-isolated cells were pelleted by centrifugation at 2,000 × g for 5 minutes at 4°C, washed in PBS, and subsequently resuspended in lysis buffer. Lysates were stored at-80°C until RNA extraction. Small RNAs were isolated following manufacturer’s instructions (illustra RNAspin Mini RNA isolation kit-GE Healthcare). 3′ adapter ligation was performed using SRBC barcode adapters specific to each sample. To monitor ligation efficiency and product size, synthetic 18-mer and 30-mer RNA markers were included. Following adapter ligation, the 3′-ligated small RNAs were size-selected on a 15% denaturing urea polyacrylamide gel, run at a constant power of 40–50 W for approximately 45-60 minutes. Gels were stained with 0.05% SYBR Gold in 0.5X TBE, and RNA fragments ranging from 18 to 30 nucleotides were excised. RNA was extracted from gel slices using the Zymo ZR Small RNA PAGE Recovery Kit, following the manufacturer’s instructions. Purified 3′-ligated RNAs were eluted in a 5′ linker mix containing the 5′ adapter. These RNAs were denatured at 70°C for 5 minutes, immediately snap-cooled on ice, and ligated to the 5′ adapter using T4 RNA ligase (NEB) and 3’ adapter using T4 RNA ligase 2 (truncated KQ. NEB) both at 16°C overnight. The resulting doubly ligated RNAs were then purified using magnetic beads (Molecular Biology Service, Vienna Biocenger). Briefly, MBS buffer and bead slurry were combined with the sample, mixed by vortexing, followed by addition of isopropanol and incubation at room temperature. Magnetic separation was used to isolate the beads; the supernatant was removed, and the beads were washed with ethanol. RNA was finally eluted in ultrapure water and transferred to PCR strips for reverse transcription. Reverse transcription was performed using a small-RNAseq RT primer specific to each sample. A no-enzyme control was included to check for contamination. First-strand cDNA synthesis was carried out using SuperScript II Reverse Transcriptase (Invitrogen). To amplify the cDNA libraries, the KAPA HiFi Real-Time Library Amplification Kit (Roche) was used. PCR reactions were set up with TruSeq Universal Adapter Forward Primer (Solexa_PCR_fwd) and TruSeq Index Reverse Primers (Solexa_IDX_rev), the latter incorporating unique sample barcodes. The PCR reaction mix included KAPA HiFi HS ReadyMix and was cycled using the following program: initial denaturation at 98°C for 45 seconds, followed by ∼12-18 cycles of 98°C for 15 seconds, 65°C for 30 seconds, and 72°C for 30 seconds, with a final extension at 72°C for 1 minute. Amplified libraries were purified using a 3% Low-Range Ultra Agarose gel (Bio-Rad) run at 80–100 V. DNA size was verified using the GeneRuler 50 bp DNA Ladder (ThermoFisher Scientific), and fragments between 150–200 bp were excised under long-wave UV light. Gel slices were transferred to clean 15 mL Falcon tubes, DNA was purified using the Zymoclean Gel DNA Recovery Kit (Zymo Research), and sequenced using 50bp single-end reads on the Illumina HiSeq V4 platform. For each developmental stage, three biological replicates were included in the analysis.

### Embryo transfection and perturbation experiments

For miRNA knockdown studies, steric blocking antisense miRNA inhibitors, that hybridize to mature miRNA species (IDT miRNA inhibitors), were diluted at 10µM in 10 mM Tris pH8.0 and injected as previously described ^26,35^. For miRNA overexpression experiments, we used chemically synthesized double-stranded RNAs that mimic mature endogenous miRNA activity (miScript miRNA mimics, Qiagen) diluted at 33µM each in 10 mM Tris pH8.0. The miRNA mimics and inhibitors were designed based on mature miRNA sequences obtained from miRbase.

### RNA Extraction, Library Preparation and Sequencing to identify miRNAs target genes

To identify to the miRNAs target genes, we performed mRNA-seq on migratory and condensing stages of the NCCs, embryos were injected and electroporated with a combination of EMT-miRNAs inhibitors with a GFP carrier (0.3 µg/µl) at stage HH5 for migratory, and MET-miRNAs inhibitors at HH8+ for condensing NCC, respectively. The embryos were then allowed to grow until they reached migratory (HH10) and condensing (HH16) stages. For the migratory stage, dorsal tubes and neural crest migration areas were dissected from the treated side with inhibitors and a control side (uninjected). For the condensing stage, trigeminal ganglia were dissected from the side treated with inhibitors and the untreated control side. Each pool was conformed with ten half embryos. Three biological replicas for each stage were used for analysis. mRNA sequencing was performed on these cell populations. The tissues were pelleted and frozen immediately at –80°C to the posterior RNA extraction. The RNA was extracted using Norgen Biotek corporation Total RNA purification kit (Cat. #17200, Norgen Biotek corporation) according to manufacturer’s instructions. DNase I RNase free (Invitrogen, Thermofisher) treatment was performed for 15 minutes at room temperature. The quality of the samples and sequencing were performed by the Macrogen company, RNA integrity was checked using 4200 TapeStation System (Agilent, Part# G2991BA), the samples with RIN>7 and rRNA Ratio≥1.5 were used. Sequencing libraries were prepared using Illumina TruSeq stranded mRNA kit and sequenced using 150bp paired-end read length on the NovaSeqX platform. Sequencing quality was assessed with FastQC (v0.11.9, ^36^). For microRNA mapping, no trimming was required, as the aligner is optimized for small RNA reads. Reads were mapped to the Gallus gallus reference genome (Galgal5) using subread-align (v2.0.1,^37^) with default parameters recommended for microRNA alignment.

Read quantification was performed with featureCounts using a microRNA annotation file from miRBase ^38^, corresponding to the Galgal5 genome assembly. Differential expression analysis was conducted using DESeq2 (v1.49.1^39^). Data visualization included volcano plots generated with EnhancedVolcano and heatmaps created using ComplexHeatmap (v2.6.2) in R.

### RNAseq Analysis

Sequence reads quality was assessed using FastQC version 0.12.1 ^36^. Adapter trimming and low-quality reads filtering was performed with TrimGalore using default parameters (v0.6.10; https://github.com/FelixKrueger/TrimGalore). The surviving reads were used as input for transcript quantification by Salmon (v0.14.1; ^40^. Quantification was performed over *G. gallus* reference transcript sequences (Galgal6a; ensembl release 106); rRNA transcripts were excluded from the analysis. The parameters used in the quantification are listed next:-l IU--discardOrphans--validateMappings. The resulting expression matrix was employed as input for the DESeq2 package (v1.46.0; ^39^. The principal component analysis (PCA) and the differential expression analysis was conducted by the DESeq2 package. All the analysis was performed in a R environment (v.4.4.2). For the Gene Set Enrichment Analysis (GSEA) results from DESeq2 analysis were used for a metric construction as follows: log2fc*-log10(pvalue). For all the detected transcripts in the set a sorted pre-ranked list was constructed based on the above-described metric. GSEA was performed using the gseGO function from the cluster profiler package ^41^. The parameters used for analysis were: minGSSize = 50, maxGSSize = 1000, pvalueCutoff = 0.05, eps = 0, pAdjustMethod = “BH”. Only ontologies for biological processes were considered. Plots were made with the enrichplot package^42^. All analysis and plots were conducted under a R environment (v4.4.2).

To identify functional relationships between upregulated genes following miRNA inhibition, protein-protein interaction networks were constructed using the STRING database (v12.0; ^43^). Genes significantly upregulated upon MET-miRNA inhibition were uploaded to the STRING online platform (https://string-db.org/), and analysis was restricted to *Gallus gallus*. The confidence interaction score was set to “medium” (0.4), and both direct (physical) and indirect (functional) associations were considered. Known interactions were derived from curated databases and experimental data, while predicted interactions included gene co-occurrence, co-expression, and protein homology. The resulting network was adapted and exported directly from STRING, with nodes corresponding to proteins and edges to predicted or validated interactions.

### Grafting experiments

The donor GFP transgenic embryos were electroporated in the dorsal neural tube with a combination of miRNA mimics targeting miR-23b-3p and miR-363-5p or with a non-targeting control mimic. Electroporations were performed at the premigratory stage, corresponding to HH8+ in the cranial region and HH9+ in the trunk. After a few hours of incubation to allow for initial expression of the miRNA mimics, electroporated neural tube explants were dissected from either the cranial (midbrain) or truncal region of the donor embryos. Concomitantly, the dorsal neural tube was ablated in the cranial region of wild-type HH9 host embryos. Donor explants were transplanted into host embryos in the same anteroposterior orientation and at equivalent axial levels to preserve regional identity. Chimeric embryos were reincubated for an additional 72 hours to allow neural crest condensation in the trigeminal ganglia. Embryos were then fixed in 4% paraformaldehyde, embedded in 7.5% gelatin 15% sucrose, and cryosectioned at 12 µm for subsequent immunohistochemistry and imaging analyses.

### Immunohistochemistry

For immunohistochemistry, embryos were fixed for 15 minutes at room temperature (RT) or overnight at 4°C in 1X phosphate buffer saline (PBS) containing 4% paraformaldehyde (PFA) and processed immediately. The embryos were washed in PBS 1X with 0.5% Triton (PBS-T) three times and subsequently blocked with 5% FBS in PBS-T for minimum 1h at RT. The primary antibodies used were anti-PAX7 IgG1 (Developmental Studies Hybridoma Bank, 1:10) and mouse anti-TUJ1 (1:200; Covance). The embryos were incubated at 4°C overnight in the primary antibody solution. Secondary antibodies used were goat anti-mouse IgG1 647 (all from Molecular Probes, 1: 500) and goat anti-mouse IgG2A Alexa Fluor 647 (1:300) overnight at 4°C. Then washed three times in PBS-T 1X and whole-mount imaged using Carl ZEISS Axio observer 7 inverted microscope (Axio observer Colibri 7, Axiocam 305 color, Axiocam 503 mono) and Carl ZEISS ZEN 2.3 (blue edition) software.

### Cryosectioning

For histological analysis, embryos were incubated in 5% sucrose for one hour at RT, and in 15% sucrose overnight at 4°C. Embryos were then incubated in 7.5% porcine gelatin for 4h at 37 °C, placed in silicone molds, snap-frozen in liquid nitrogen, and stored at −80 °C. Transverse sections of 10-15um were obtained for cryosectioning (provided by Lic. Gabriela Carina López from the “Histotechnical Service” at INTECH). After slides were washed in PBS with 0.5% Triton (PBSt) at 42 °C for 10 min for gelatin removal, rinsed twice in PBS, and mounted with Fluoromount (ThermoFisher Scientific) Mounting Medium for imaging.

### Quantification of neural crest migration area and cell number

To measure the area of migratory neural crest cells, whole-mount embryos stained for PAX7 were imaged under identical settings. Images were processed in FIJI (ImageJ v1.53; ^44^), and background fluorescence was subtracted using reference regions due to uniform background levels. Using the line tool, a region of interest (ROI) was manually drawn around the area of migratory PAX7+ cells on both the electroporated and contralateral control side. The area on the treated side was normalized to the internal control side to calculate the relative migratory area per embryo.

To quantify the contribution of neural crest cells in transplantation experiments, GFP+ neural crest-derived neurons expressing Tuj1+ within the trigeminal ganglion (TG) were counted using the cell counter tool in FIJI. Cell counts were compared between treatment groups, including control mimics from both cranial (Cra/Cra) and trunk (Trunk/Cra) sources and MET-miRNA mimic treated trunk transplants.

Statistical analyses were performed using GraphPad Prism v9. Comparisons were evaluated using two-tailed unpaired t-tests or one-way ANOVA followed by post hoc tests, as appropriate. Statistical significance was defined as P < 0.05.

## RESULTS

### Transcriptional miRNA profiling during cranial neural crest transitions

To identify the key up-regulated miRNAs involved in regulating NCC delamination (EMT-miRNAs) and condensation (MET-miRNAs), we performed miRNA sequencing on pure populations of NCCs at three different developmental stages **(Fig. 1A)**. We electroporated the neural tube with cranial NCC-specific enhancers, NC1 or 10E2, driving Citrine expression in pre-migratory/early migratory or late-migratory cells, respectively ^34,45^. Fluorescence-activated cell sorting was then used to isolate Citrine+ NCCs and surrounding Citrine-non-NCC cells from chicken embryos at pre-migratory (Hamburger and Hamilton stage 9, HH9), migratory (HH10-11), and condensing (HH16) stages. The results of Principal Component Analysis (PCA) showed that the RNAseq samples were well correlated, clearly separating the 3 timepoints **(Fig. S1B)**. Further hierarchical clustering of miRNA expression clustered all detected miRNAs into 10 different groups **(Fig. S1A, C)**.

**Figure 1.**
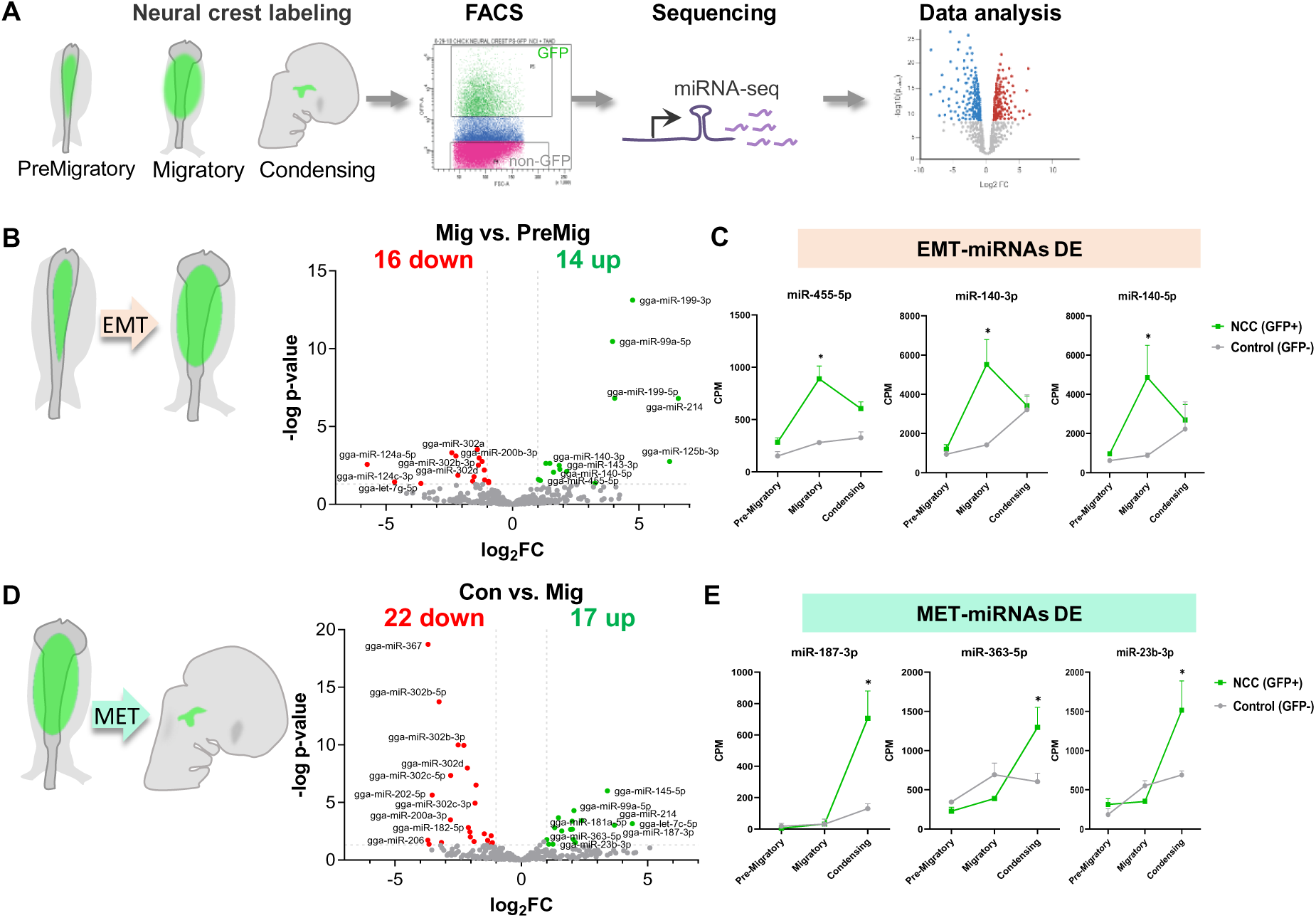
Stage-specific microRNA profiling reveals EMT-and MET-associated miRNAs in neural crest cells. **(A)** Schematic of the experimental workflow for microRNA sequencing (miRNA-seq) of neural crest cells (NCCs) at distinct developmental stages in chick embryos: PreMigratory (5–6 somite stage), Migratory (8–10ss), and Condensing (16ss). Citrine+ NCCs were labeled using NC1 and Sox10E2 enhancers and isolated via fluorescence-activated cell sorting (FACS) into GFP+ (NCC) and GFP– (non-NCC) populations. miRNA libraries were prepared from each population and analyzed by differential expression. **(B, D)** Volcano plots showing differentially expressed miRNAs between Migratory vs. PreMigratory stages **(B)** and Condensing vs. Migratory stages **(D)**. Significantly downregulated (red) and upregulated (green) miRNAs are highlighted (cutoff: |log₂ FC| > 1, *p* < 0.05). **(C, E)** Expression profiles (CPM) of representative miRNAs associated with epithelial-to-mesenchymal transition (EMT; C) or mesenchymal-to-epithelial transition (MET; E), showing stage-specific enrichment in NCCs (GFP+) compared to non-NCC controls (GFP–). Data represent mean ± SEM; \**p* < 0.05.

Differential miRNA expression analysis between migratory and pre-migratory stages revealed 14 significantly enriched EMT-miRNAs **(Fig. 1B)**, with only three (miR-140-3p, miR-140-5p, and miR-455-5p) exclusively detected in NCCs **(Fig. 1C, S2, S4, Table S1)**. Similarly, a comparison between condensing and migratory stages identified 17 significantly enriched MET-miRNAs **(Fig. 1D)**, with three (miR-23b-3p, miR-187-3p, and miR-363-5p) detected only in NCCs **(Fig. 1E, S3, S4, Table S1)**.

**Figure 2.**
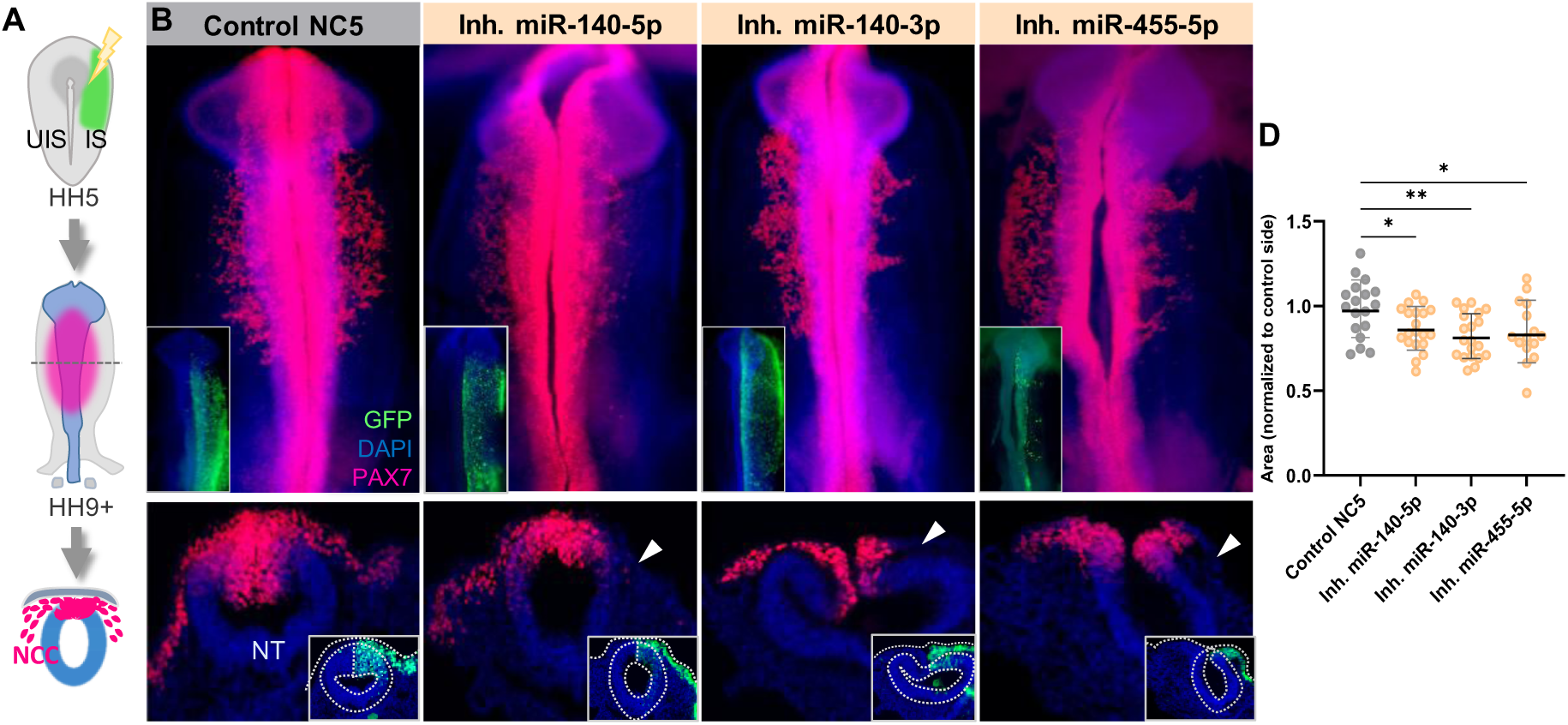
Inhibition of EMT-associated miRNAs disrupts neural crest cell migration. **(A)** Schematic of the experimental design. Chick embryos were electroporated at gastrula stages (HH5) with miRNA inhibitors and a GFP reporter on the right side (experimental), and allowed to develop to HH9+, when cranial neural crest cells (NCCs) have migrated. **(B)** Representative images of whole-mount and transverse sections of embryos stained for PAX7 (magenta) and DAPI (blue) to assess NCC migration. Insets show the GFP+ (green) electroporated side. Compared to controls (NC5 inhibitor), embryos treated with inhibitors for miR-140-5p, miR-140-3p, or miR-455-5p exhibit reduced NCC migration (arrowheads in transverse sections). NT, neural tube. **(C)** Quantification of the area occupied by migrating PAX7+ cells on the electroporated side, normalized to the contralateral control side. Each dot represents one embryo. Bars indicate mean ± SEM; \**p* < 0.05, \*\**p* < 0.01.

**Figure 3.**
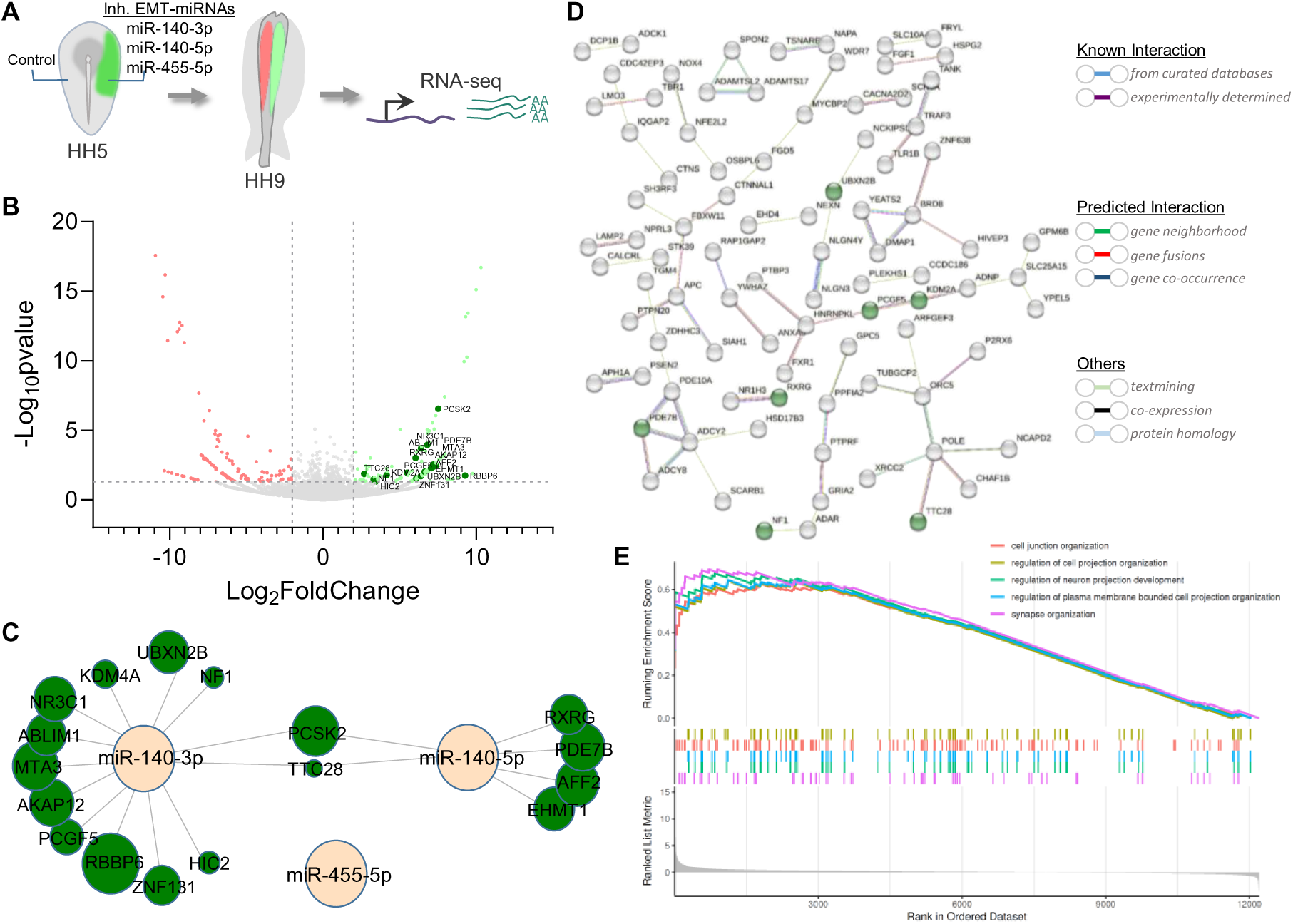
Transcriptomic analysis identifies putative target genes regulated by EMT-associated miRNAs. **(A)** Experimental overview. Chick embryos were electroporated on the right side at HH5 with inhibitors for EMT-associated miRNAs (miR-140-3p, miR-140-5p, and miR-455-5p), and tissues were collected at HH9 for RNA-seq. The left, untreated side served as an internal control. **(B)** Volcano plot of differentially expressed genes following EMT-miRNA inhibition. Upregulated genes predicted as direct miRNA targets are highlighted in dark green (|log₂ FC| > 1, p < 0.05). **(C)** miRNA–mRNA interaction network linking EMT-miRNAs (peach nodes) to upregulated predicted targets (green nodes). Node size reflects fold change values from RNA-seq. **(D)** STRING-based protein-protein interaction network of the upregulated genes (gray nodes) and predicted EMT-miRNA targets (green nodes). Edges indicate interactions, with edge type indicated in the legend. **(E)** Gene Set Enrichment Analysis (GSEA) for the EMT-miRNA inhibition transcriptomic data set. Top five enriched GO terms are shown with enrichment scores and ranked gene distributions.

**Figure 4.**
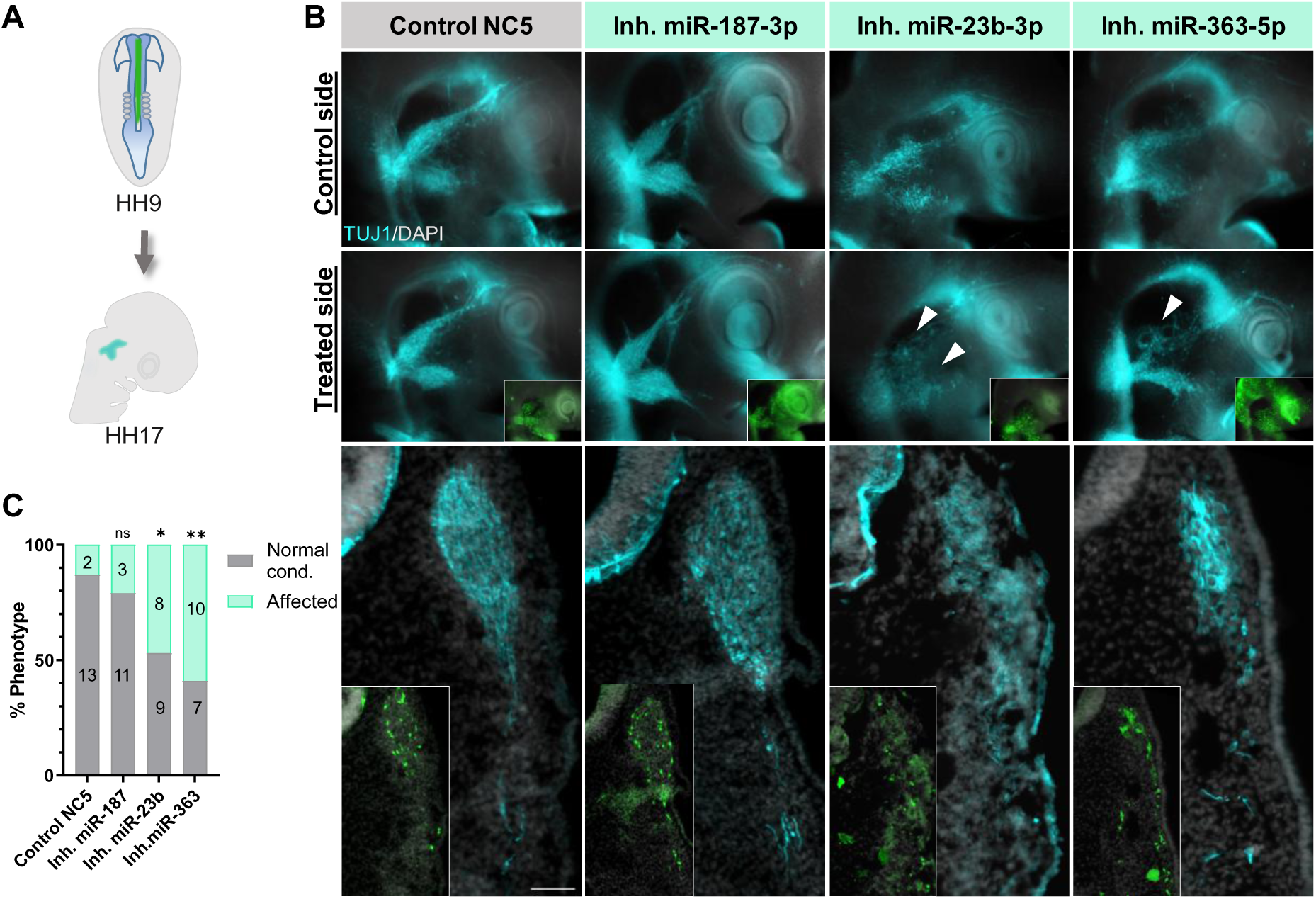
Inhibition of MET-associated miRNAs impairs trigeminal ganglion morphogenesis. **(A)** Schematic of the experimental design. Chick embryos were electroporated at HH9 on the right side with inhibitors targeting MET-associated miRNAs (miR-187-3p, miR-23b-3p, miR-363-5p) and allowed to develop until the condensation stage (HH17). **(B)** Representative images of whole-mount (top control and treated side panels) and transverse sections (bottom panels) of embryos stained for TUJ1 (cyan) and DAPI (blue) to assess trigeminal ganglion morphology. Insets show the GFP+ (green) electroporated treated side. Compared to controls (NC5 inhibitor), embryos treated with inhibitors for miR-23b-3p or miR-363-5p, but not for miR-187-3p, exhibit disrupted trigeminal ganglion condensation or nerve branching (white arrowheads), respectively. **(C)** Quantification of phenotypic outcomes, showing the number embryos having affected (light green) or normal (gray) trigeminal ganglion morphology compared to controls. Statistical analysis was performed using a χ² test (\**p* < 0.05, \*\**p* < 0.01, ns = not significant).

### Loss of EMT-miRNAs causes defects in NCC migration

As an initial step in assessing the role of core EMT-miRNAs for NCC delamination, we individually inhibited miR-140-5p, miR-140-3p, and miR-455-5p (Sequence in **Table S2**). Specific antisense miRNA inhibitors were unilaterally injected and electroporated into gastrula stage embryos (HH5), which were then grown until stage HH9+ **(Fig. 2A)**. This treatment led to an increase in the number of embryos exhibiting NCC migration defects, as indicated by PAX7 staining on the injected side compared to the uninjected side within the same group of embryos, or embryos injected with control inhibitor (Control NC5) **(Fig. 2B)**. The area measurement of migratory PAX7+ cells revealed a significant reduction in the injected side (IS), normalized to the control uninjected side (UIS), compared to control NC5-treated embryos **(Fig. 2C)**. These results indicate that the identified EMT-miRNAs are required for proper NCC delamination.

### Treatment with EMT-miRNA inhibitors alters the expression of potential target genes

To identify potential targets of EMT-miRNAs, we injected gastrula-stage embryos with a combination of three miRNA inhibitors for (Inh. EMT-miRNAs) on the right side and allowed them to develop until the stage of delamination (HH9+). We then dissected the treated and control side tissues and performed bulk RNA-seq (**Fig. 3A)**. Differential expression analysis identified 251 transcripts with significant expression changes (log2FC > 2) out of the 37,542 identified **(Fig. 3B, Table S4)**. Since miRNAs generally act to suppress the expression of their target genes, we focused our analysis on the upregulated transcripts. To determine whether any of the upregulated genes are predicted target of the EMT-miRNAs, we performed an *in silico* analysis using TargetScan and miRDataBase (full list of predicted targets is in **table S3**). This analysis identified 17 upregulated genes as predicted targets of miR-140-5p and miR-140-3p, but not for miR-455-5p **(Fig. 3B-C)**. Notably, several of these predicted targets, highlighted in green, form functional protein association networks with other upregulated genes, as revealed by STRING analysis^43^ **(Fig. 3D)**. Importantly, several of the target genes are epigenetic regulators (KDM4A^46^, MTA3, PCGF5^47^, EHMT1), transcriptional regulators (NF1^48^, NR3C1, ZNF131, HIC2, RXRG, AFF2) or involved in cell adhesion/migration (ABLIM1^49^, AKAP12^50^). Therefore, we next performed a gene set enrichment analysis (GSEA,^51^) to further explore the transcriptomic dataset. The top five enrichment categories among the data set were associated with cell junction organization, regulation of cell projections, organization of plasma membrane-bounded projections, and synapse organization **(Fig. 3E, Table S5)**.

Together, these findings identify putative target genes that are likely repressed by EMT-miRNAs and may contribute to the transcriptional regulation required for proper NCC delamination and migration.

### Loss of MET-miRNAs alters trigeminal ganglion morphogenesis

To assess the requirement of the identified MET-miRNAs during NCC condensation and trigeminal ganglion formation, we inhibited miR-23b-3p, miR-187-3p, and miR-363-5p (Sequence in **Table S2)** by electroporating specific inhibitors during the pre-migratory NCC stage. The experimental strategy is outlined in **figure 4A**: the right neural tube of HH9 embryos was electroporated with the respective miRNA inhibitors, and the embryos were subsequently analyzed at HH17 by IHC for Tuj1 to evaluate ganglion formation. Our results showed that inhibition of miR-23b-3p led to reduced condensation of the ganglion on the treated side, as indicated by arrowheads **(Fig. 4B)**. In contrast, inhibition of miR-363-5p caused pronounced disorganization, particularly affecting the branching of the nerves from the ophthalmic lobe. Inhibition of miR-187-3p, however, produced minimal disruption, resembling the control side and embryos treated with the Control NC5 inhibitor. Quantification of these phenotypes, showing the percentage of embryos with normal versus affected trigeminal ganglion formation for each condition, is summarized in **figure 4C**.

These findings demonstrate that miRNAs associated with the mesenchymal-to-epithelial transition, particularly miR-23b and miR-363, are critical for proper NCC condensation and trigeminal ganglion formation.

### Inhibition of MET-miRNAs reveals their potential target genes

To identify potential target genes of MET-miRNAs, we performed RNA-seq on trigeminal ganglia from embryos electroporated with inhibitors against the three MET-miRNA (miR-23b-3p, miR-363-5p and miR-187-3p). We injected the inhibitor into the neural tube at stage HH9 and electroporated the right side of the embryos, allowing them to develop until HH17. Trigeminal ganglia were then dissected from both the treated and control sides, pooling 10 ganglia per condition from three independent electroporations for RNA-seq analysis **(Fig. 5A)**. Differential expression analysis revealed significant transcriptional changes following MET-miRNA inhibition. Out of all the analyzed transcripts, 289 showed differential expression (log2FC > 2, p-value < 0.05, **Table S4**). Notably, seven of the upregulated genes were predicted targets, identified by *in silico* analysis using TargetScan and miRDB (full list of predicted targets is in **table S3**), particularly those of miR-23b-3p and miR-363-5p **(Fig. 5B-C)**. Consistent with the absence of a discernible phenotype following miR-187-3p inhibition, none of the upregulated genes were predicted targets of this miRNA. Notably, several of these predicted targets, highlighted in green, form functional protein association networks with other upregulated genes, as revealed by STRING analysis **(Fig. 5D)**. Importantly, several of the target genes are epigenetic regulators (BRWD1, ^52^), transcriptional regulators (ZBTB18, ^53^), regulators of vesicle trafficking (TMEM131L ^54^, SYTL3^55^) and cytoskeleton remodeling factors (MAP4K4^56^, CTTNBP2NL). We next used GSEA to further analyze the transcriptomic dataset. The top five enrichment categories were associated with aerobic and cellular respiration, energy derivation by oxidation of organic compounds, generation of precursor metabolites and energy, and translation **(Fig. 5E, Table S5)**.

**Figure 5.**
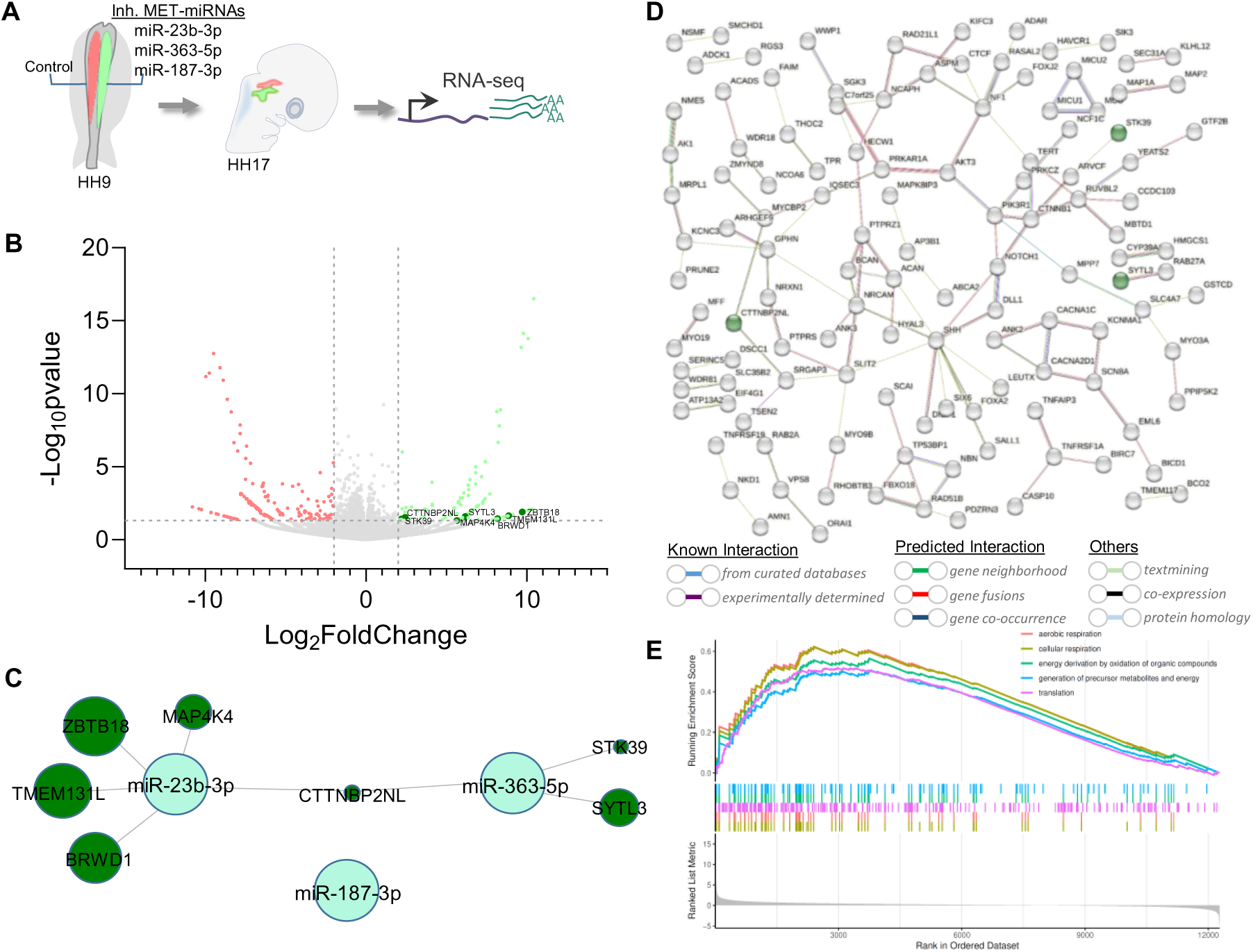
Transcriptomic analysis identifies putative target genes regulated by MET-associated miRNAs. **(A)** Schematic of the experimental design. Chick embryos were electroporated at HH9 on the right side with inhibitors targeting MET-associated miRNAs (miR-23b-3p, miR-363-5p, and miR-187-3p) and allowed to develop until the trigeminal ganglion condensation stage (HH17). Treated and untreated (control) tissues were then dissected separately for RNA sequencing. **(B)** Volcano plot showing differentially expressed genes following MET-miRNA inhibitor treatment. Upregulated genes predicted as MET-miRNA targets are highlighted in dark green. **(C)** miRNA–mRNA interaction network linking MET-miRNAs (light green nodes) to upregulated predicted targets (green nodes). Node size reflects fold change values from RNA-seq. **(D)** STRING protein–protein interaction network for upregulated genes, illustrating known and predicted functional associations among candidate MET-miRNA targets. **(E)** Gene Set Enrichment Analysis (GSEA) for the MET-miRNA inhibition transcriptomic data set. Top five enriched GO terms are shown with enrichment scores and ranked gene distributions.

Together, these findings identify putative target genes that are likely repressed by MET-miRNAs and may contribute to the transcriptional regulation required for proper NCC condensation and/or trigeminal ganglion morphogenesis.

### Introduction of MET-miRNAs in trunk NCC enhances their ability to condensate and contribute into neurons of the trigeminal ganglia

Previously elegant quail-chick chimeric grafting experiments of cardiac or trunk NCC transplanted into the midbrain region demonstrated that such grafts yielded trigeminal ganglia that were smaller and contained fewer neurons compared to control midbrain-derived NCC grafts ^3^. These results suggest that there are differences in their ability to condense and differentiate into trigeminal neurons in an axial level selective manner.

To test whether MET-associated microRNAs (MET-miRNAs) contribute to these properties, we electroporated MET-miRNA mimics (miR-23b and miR-363) or control mimics into the neural tubes of GFP-transgenic chick embryos at stage HH8+. Dorsal neural tubes were then dissected from cranial (HH9) or trunk (HH10) levels and grafted into wild-type host embryos at stage HH9. Chimeric embryos were cultured until stage HH17 and subsequently analyzed **(Fig. 6A)**. As expected, control mimic cranial-to-cranial chimeras displayed robust condensation of GFP+ NCCs within the TG **(Fig. 6B-C)**. Consistent with prior findings, grafting trunk NCCs electroporated with control mimics into the midbrain region resulted in significantly fewer GFP+ cells integrating into the TG. Notably, trunk NCCs overexpressing MET-miRNAs exhibited a markedly enhanced contribution to TG formation, with GFP+ cell numbers approaching those observed in cranial-to-cranial grafts.

**Figure 6.**
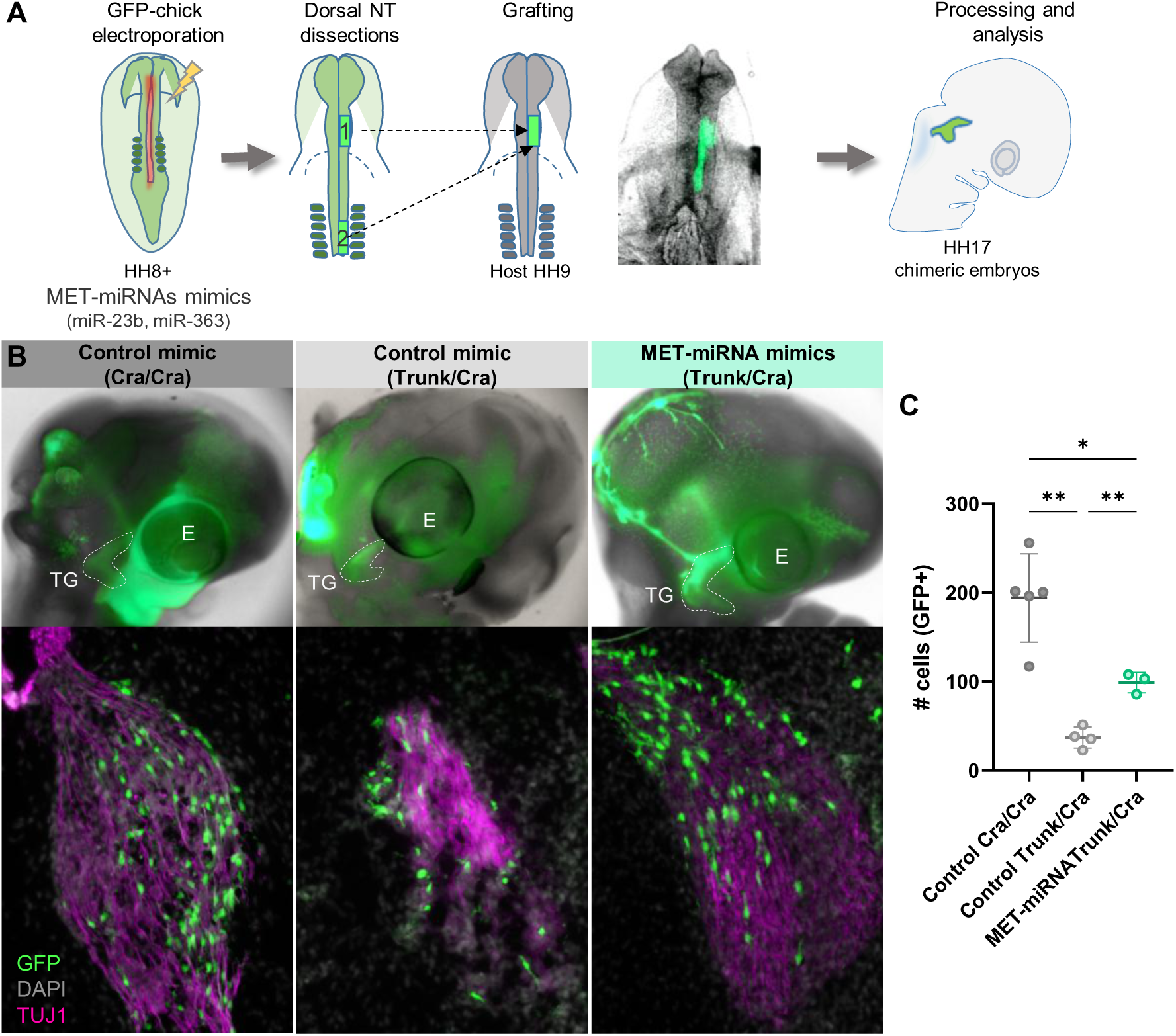
Ectopic expression of MET-associated miRNAs enhances the ability of trunk NCCs to contribute to trigeminal ganglion neurons. **(A)** Schematic of the experimental design. Transgenic GFP-chick embryos were electroporated at HH8+ with Control or MET-associated miRNA mimics (miR-23b-3p and miR-363-5p). Dorsal neural tube explants from trunk or cranial regions were then dissected and grafted into the cranial region of host embryos at HH9. Chimeric embryos were collected at HH17 for analysis. **(B)** Representative whole-mount and transverse section images of chimeric embryos showing GFP+ cells (green) within the trigeminal ganglion (TG) stained for TUJ1 (magenta) and DAPI (gray). Control mimics from both cranial (Cra/Cra) and trunk (Trunk/Cra) sources were compared to MET-miRNA mimic-treated trunk grafts. **(C)** Quantification of GFP+ cells within the TG. Trunk NCCs expressing MET-miRNA mimics contributed significantly more to differentiated neurons than control trunk NCCs. Statistical significance was assessed using one-way ANOVA with post hoc tests (*p < 0.05, **p < 0.01).

These results indicate that MET-miRNAs enhance the capacity of NCCs to contribute to TG formation, supporting a role for miRNA-mediated regulation in promoting NCC delamination and condensation—processes critical for proper cranial gangliogenesis.

## DISCUSSION

Cranial NCCs undergo a tightly regulated sequence of EMT and MET-like events that are critical for their migration and subsequent condensation into structures such as the trigeminal ganglion. Given the central role of these transitions in NCC behavior coupled with their known involvement in tumor metastasis, we examined the possible contribution of miRNAs to these processes, as they are well-established regulators of EMT and MET in cancer and tissue morphogenesis ^57^. To this end, we performed miRNA transcriptomic analysis across distinct stages of NCC development. We identified differentially expressed miRNAs between the premigratory and migratory stages (EMT), as well as between migratory and condensing stages (MET). Several of these miRNAs were also differentially expressed in NCCs relative to non-NCC control populations. This analysis led to the identification of three EMT-associated miRNAs (miR-140-3p, miR-140-5p, miR-455-5p) and three MET-associated miRNAs (miR-23b-3p, miR-187-3p, miR-363-5p).

We focused our investigation on miRNAs with dynamic expression changes at the onset of NCC migration and condensation. Among the EMT-miRNAs, miR-140-3p is known to be involved in cartilage development and NCC-driven craniofacial morphogenesis. Consistent with our findings, changes in miR-140-3p levels alter the expression of its direct target *Pdgfra*, a gene essential for proper NCC dispersion and palatogenesis in zebrafish ^58^. MiR-140-5p, another EMT-miRNA, has been shown to suppress proliferation and migration in many cancers, including Wilms’ tumor, hepatocellular carcinoma, and lung cancer ^59–62^. Importantly, and consistent with our results, downregulation of miR-140-5p in neural crest-derived SH-SY5Y cells (a model for Hirschsprung’s disease) significantly impairs both migration and proliferation ^63^. Together with the present results, these findings suggest a divergent role for miR-140-5p in NCC development compared to cancer contexts, supporting its function as a positive regulator of NCC migration and its possible involvement in the pathogenesis of neural crest-related disorders such as Hirschsprung’s disease. The third EMT-associated miRNA we identified, miR-455-5p, has been extensively linked to increased invasiveness and migratory capacity in various cancers ^64–67^. However, to our knowledge, this is the first study to associate miR-455-5p with the regulation of NCC migration *in vivo*, highlighting a potential conserved role for this miRNA in promoting cellular motility across developmental and pathological contexts.

Our RNA-seq analysis aimed at identifying potential targets of EMT-miRNAs revealed 17 genes significantly upregulated and predicted to interact with miR-140-3p (11 genes), miR-140-5p (4 genes), or both (2 genes). Notably, none of the upregulated genes were predicted targets of miR-455-5p, suggesting that this miRNA may primarily function by modulating gene expression post-transcriptionally, possibly through translational repression rather than mRNA degradation. Among the identified targets, several genes encode epigenetic regulators (e.g., KDM4A, MTA3, PCGF5, EHMT1), transcription factors and co-regulators (NF1, NR3C1, ZNF131, HIC2, RXRG, AFF2), and molecules involved in cell adhesion and migration (ABLIM1, AKAP12). It is noteworthy that some of these genes were previously implicated in the negative regulation of cell migration, including NCC. Among those, *Kdm4a* (or *JmjD2A*) encodes a histone demethylase playing an important role for NCC specification whose expression get reduced at migratory stages ^68^. *Nf1* (Neurofibromin 1) encodes a Ras GTPase-activating protein that negatively regulates Ras signaling, a pathway essential for cell motility. Genetic disruption of NF1 in mouse models has been shown to cause developmental abnormalities in several neural crest-derived tissues, including the heart and craniofacial structures. These phenotypes suggest that *Nf1* may play a regulatory role in neural crest development, potentially by modulating the balance between proliferation and migration of NCCs ^69^. Similarly, TTC28, another predicted miR-140 target, is highly expressed during craniofacial development, and its misregulation is associated with cleft palate defects ^70^.

It is important to note that several target genes of EMT-related miRNAs form functional protein association networks with other upregulated genes involved in enriched transcriptional pathways, including those related to cell junction organization, cell projection organization, and plasma membrane–bounded cell projections, among others. These findings suggest that, beyond the regulation of direct miRNA targets, EMT-miRNAs may also influence additional, indirect pathways that could impact NCC delamination and migration. Together, identification of these genes provides insight into how EMT-miRNAs may influence NCC behavior by targeting molecular pathways that coordinate epigenetic remodeling, delamination, migration, and lineage specification.

Among the MET-associated miRNAs and consistent with our findings, miR-23b was previously reported to be highly expressed in the trigeminal ganglion of mice ^23^. However, its specific role in the development of this structure has not been investigated. miR-23b-3p has been widely studied in the context of cancer, where it is generally regarded as a tumor suppressor, as its inhibition is known to promote epithelial-to-mesenchymal transition ^71–73^. Our analysis reveals that miR-23b is essential for the proper condensation of the trigeminal ganglion. Notably, to date, no studies in cancer models have demonstrated a role for miR-23b in mesenchymal-to-epithelial transition processes.

MiR-363 has been shown to suppress cell proliferation, migration, and invasion in colorectal cancer cells^70^, underscoring its potential role in regulating cell cycle and differentiation. Notably, during postnatal development in mice, miR-363-5p promotes Schwann cell myelination, while its downregulation enhances Schwann cell proliferation and migration during nerve regeneration following injury ^74^. Additionally, miR-363-5p has been implicated in promoting neurite outgrowth in dorsal root ganglion neurons ^75^. These findings align with our results, which show that inhibiting miR-363-5p impairs proper nerve outgrowth from the trigeminal ganglion.

Our RNA-seq analysis aimed to identify potential targets of MET-miRNAs revealed 7 genes significantly upregulated and predicted to interact with miR-23b-3p (4 genes), miR-363-5p (2 genes), or both (1 gene). Among them, we identified epigenetic regulators (BRWD1), transcriptional regulators (ZBTB18), regulators of vesicle trafficking (TMEM131L, SYTL3) and cytoskeleton remodeling factors (MAP4K4, CTTNBP2NL). Furthermore, target genes of MET-related miRNAs form functional protein association networks with other upregulated genes involved in enriched transcriptional pathways related to aerobic respiration and metabolic processes. Consistent with this, aerobic glycolysis, a hallmark of cancer cells, is also utilized to regulate epithelial-mesenchymal plasticity under physiological conditions ^76^. Bhattacharya et al. ^77^ demonstrated that NCC relies on transient glycolytic upregulation prior to delamination and during migration, which diminishes upon differentiation. In this context, our results position MET-miRNAs as metabolic regulators, downregulating glycolytic genes to initiate NCC condensation and trigeminal ganglion morphogenesis and differentiation.

NCC-derived neurons in the TG originate from rhombomeres r1/r2 and part of r3. Lwigale *et al.* ^3^ showed that replacing cranial NCCs with trunk NCCs reduces their contribution to TG neurons. By expressing MET-miRNAs in trunk NCCs and transplanting them into the cranial region, we demonstrated that intrinsic miRNA composition influences NCC condensation and differentiation capacities.

Strikingly, all the identified EMT and MET miRNAs appear to be vertebrate-specific, with no orthologues found in non-vertebrate chordates such as ascidians or amphioxus, nor in any other non-chordate taxon ^78^. This observation suggests that the evolutionary acquisition of novel miRNA families in vertebrates coincides with the emergence of key vertebrate innovations—most notably the neural crest—and reflects the increasing need for fine-tuned regulation of complex gene regulatory networks during their development.

## Acknowledgments

We thank all the authors and members in the Laboratory of Developmental Biology at the INTECH for their contribution and helpful discussions during the course of our study. We thank Agustina Ganuza (Associate Technician. CONICET) for her technical assistance in the lab. We are very grateful to the directors and students from “Escuela de Educación Secundaria Agraria de Chascomús” for providing fertilized eggs of excellent quality. This work was supported by the Wellcome Trust (304976_Z_24_Z to P.H.S-M.).

## Author contributions

R.B.M. and P.H.S-M. designed, performed the experiments and wrote the manuscript with editing and input from all coauthors; E.S-V, Y.E.B, E.M.S performed experiments; A.M.A contributed in the bioinformatics analyses; P.L. contributed in the grafting experiments; P.L. L.C and M.E.B contributed in the discussion, experimental design and manuscript writing.

---

**Supplementary Figure S1.**
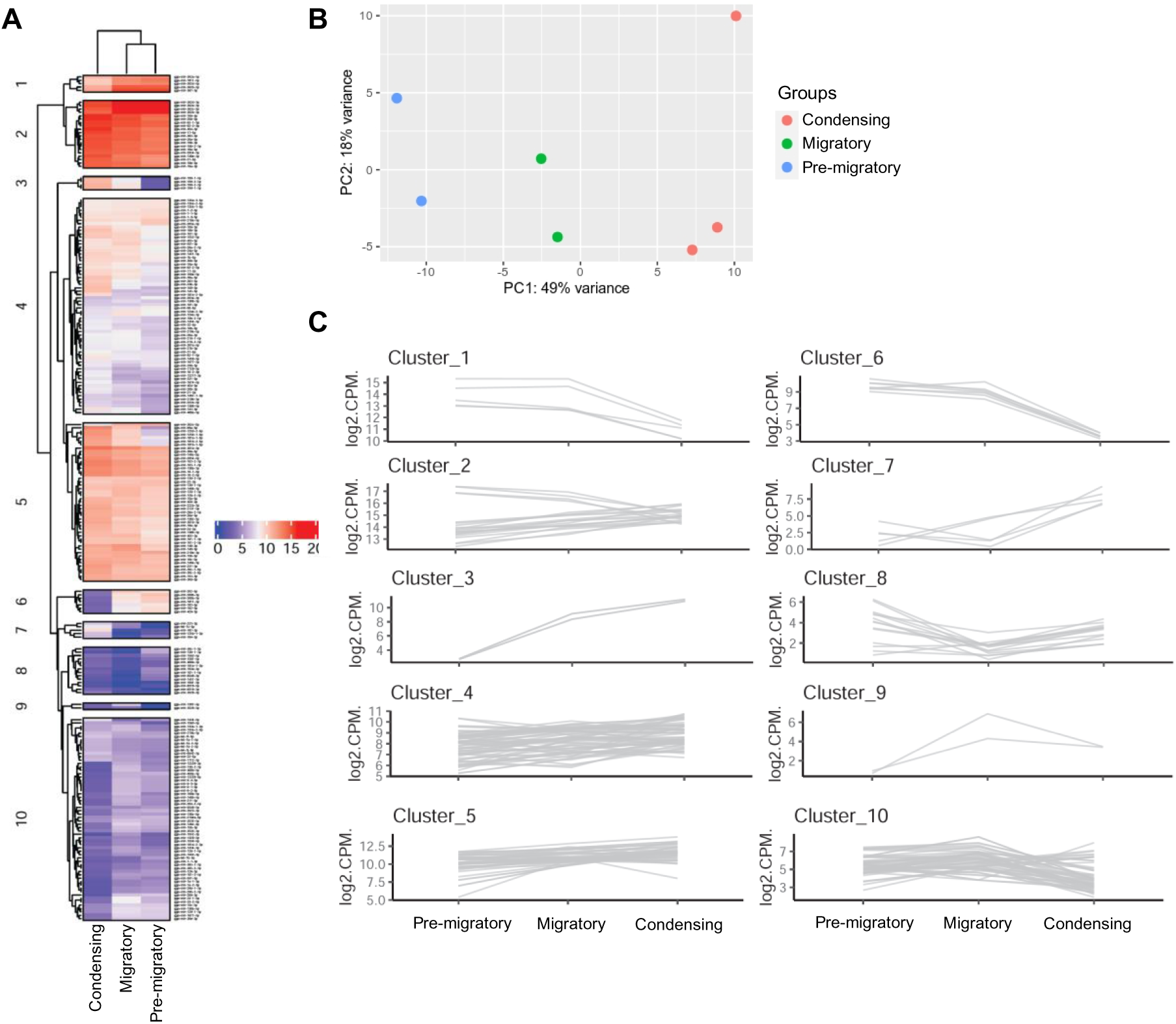
Transcriptomic profiling reveals stage-specific gene expression dynamics during trigeminal ganglion development. **(A)** Heatmap showing the hierarchical clustering of differentially expressed genes across three developmental stages of chick cranial neural crest-derived trigeminal ganglion: pre-migratory (HH9), migratory (HH12), and condensing (HH17). Ten distinct gene expression clusters were identified based on expression patterns. **(B)** Principal component analysis (PCA) of transcriptomic data showing clear separation of the three developmental stages. PC1 accounts for 49% and PC2 for 18% of the variance in the data. **(C)** Line plots representing gene expression trajectories (log₂ CPM) for each of the 10 clusters across the three developmental stages. Each gray line represents a single gene within the cluster, highlighting temporal expression dynamics during neural crest progression and trigeminal ganglion morphogenesis.

**Supplementary Figure S2.**
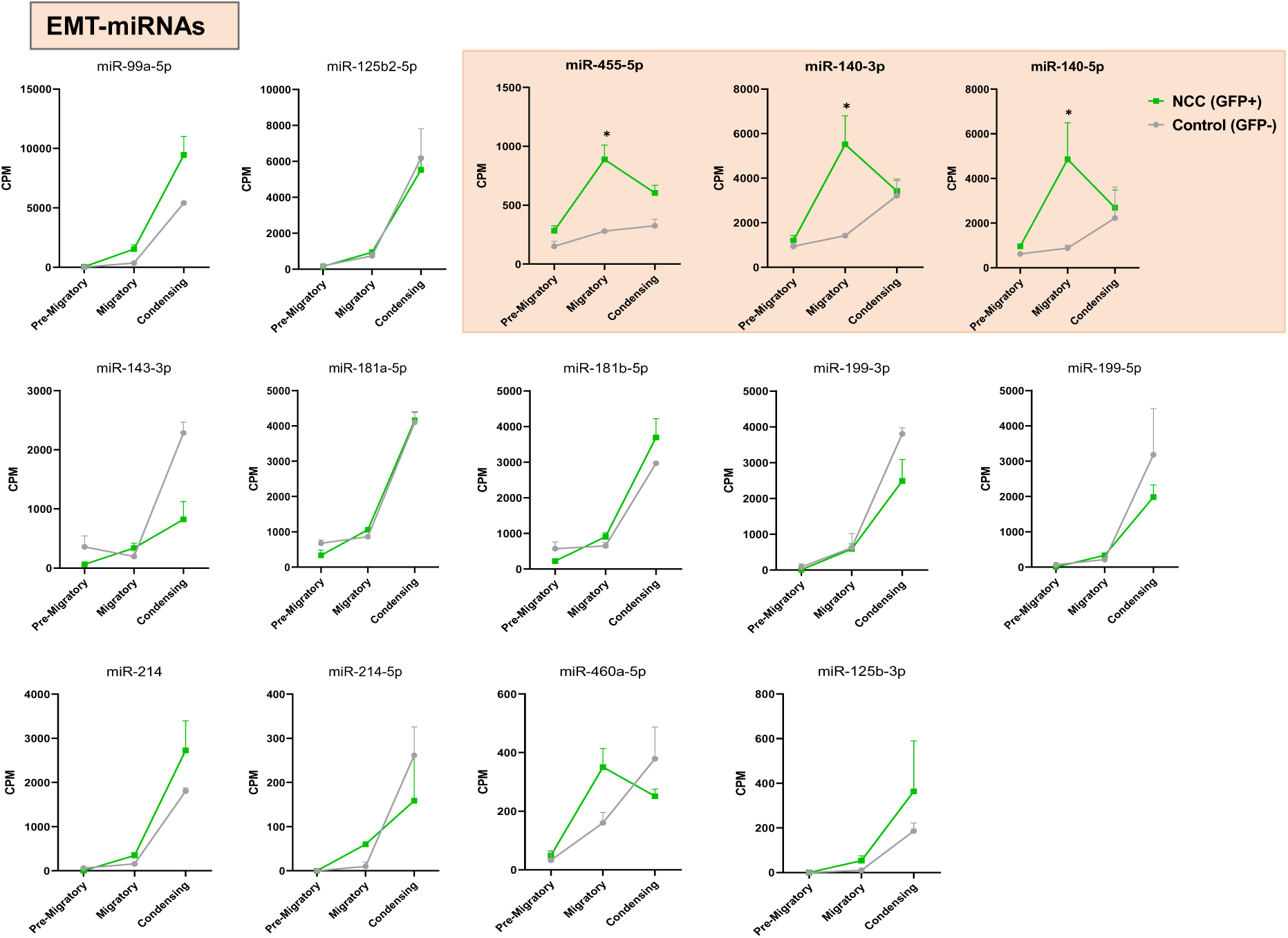
Expression dynamics of EMT-miRNAs across neural crest developmental stages. Line plots show the expression (CPM, counts per million) of EMT-miRNAs in fluorescence-activated sorted GFP⁺ (neural crest cells) and GFP⁻ (non-neural crest control) populations at pre-migratory, migratory, and condensing stages. Green lines represent GFP⁺ cells, while gray lines represent GFP⁻ cells. miRNAs highlighted in the shaded panel (miR-455-5p, miR-140-3p, miR-140-5p) show significant enrichment at the migratory stage in GFP⁺ cells (*p < 0.05), consistent with a role in EMT regulation. Data are presented as mean ± SEM.

**Supplementary Figure S3.**
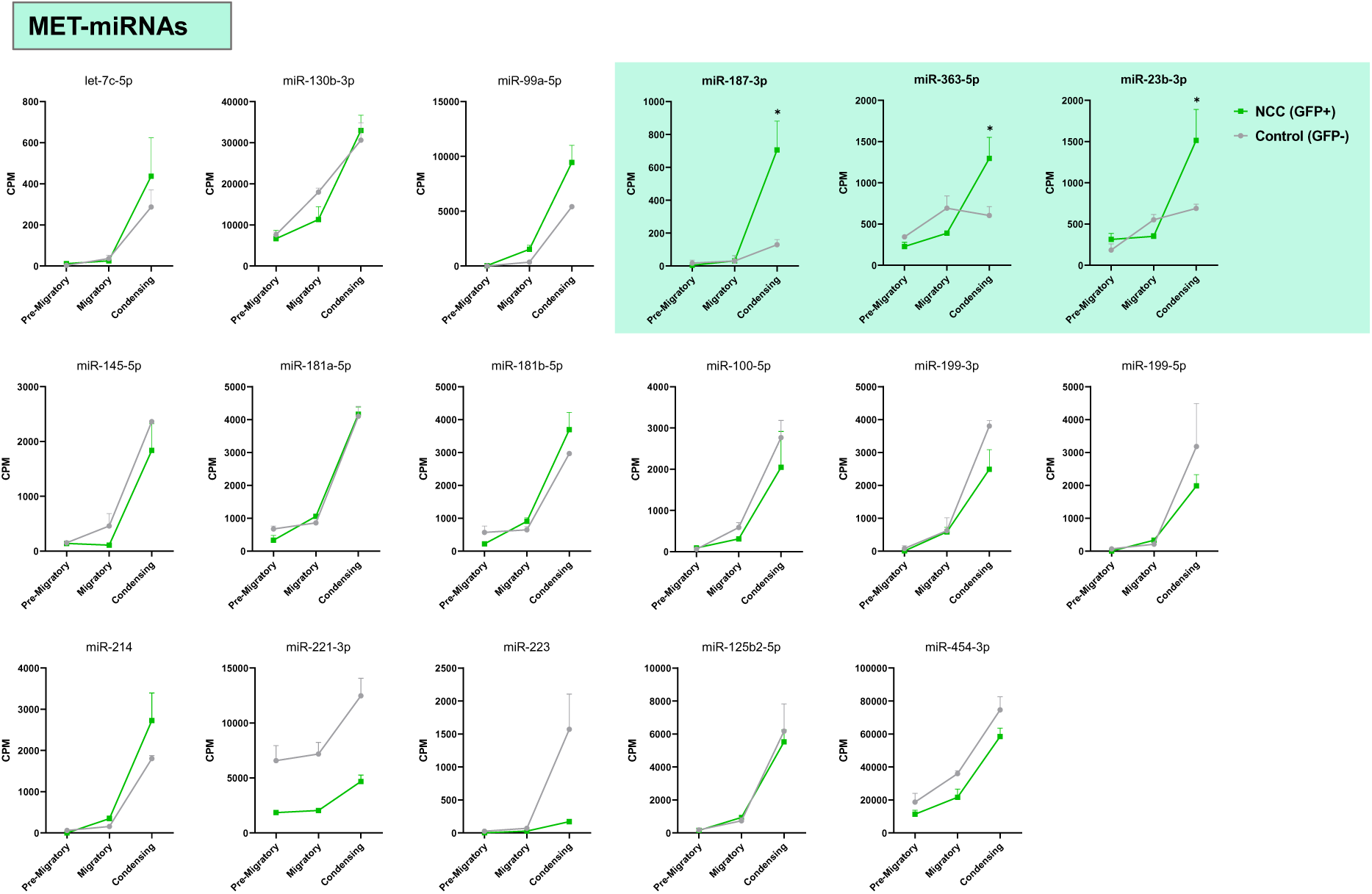
Expression dynamics of MET-miRNAs across neural crest developmental stages. Line plots show the expression (CPM, counts per million) of MET-miRNAs in fluorescence-activated sorted GFP⁺ (neural crest cells) and GFP⁻ (non-neural crest control) populations at pre-migratory, migratory, and condensing stages. Green lines represent GFP⁺ cells, while gray lines represent GFP⁻ cells. miRNAs highlighted in the shaded panel (miR-187-3p, miR-363-5p, miR-23b-3p) show significant enrichment at the condensing stage in GFP⁺ cells (*p < 0.05), consistent with a role in MET regulation. Data are presented as mean ± SEM.

**Supplementary Figure S4.**
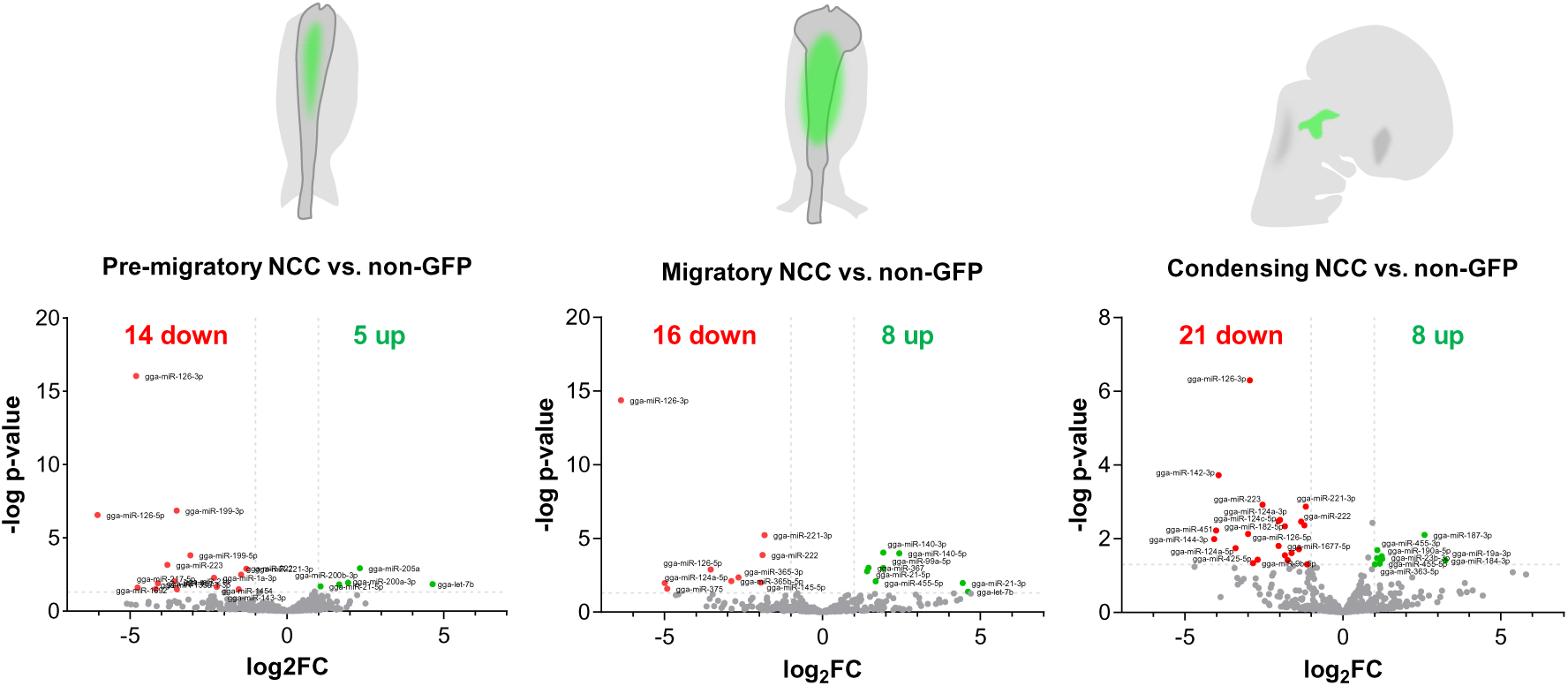
Differential miRNA expression in cranial NCCs versus surrounding non-GFP cells across developmental stages. Volcano plots show differentially expressed miRNAs between GFP⁺ neural crest cells (NCCs) and GFP⁻ surrounding cells at pre-migratory (left), migratory (middle), and condensing (right) stages. Each dot represents a miRNA, with green indicating significantly upregulated and red indicating significantly downregulated miRNAs in NCCs (cutoff: |log₂ FC| > 1, *p* < 0.05).

## SUPPLEMENTARY TABLES LEYENDS

**Table S1:** miRNA-seq data.

**Table S2:** List of utilized oligos.

**Table S3:** Lists of EMT-and MET-miRNAs target genes analyzed by TargetScan and miRDBase.

**Table S4:** RNA-seq data from embryos treated with EMT-and MET-miRNA inhibitors.

**Table S5:** GSEA analysis.

